# A comparison of wild boar and domestic pig microbiota does not reveal a loss of microbial species but an increase in alpha diversity and opportunistic genera in domestic pigs

**DOI:** 10.1101/2024.03.29.587377

**Authors:** Rajibur Rahman, Janelle M. Fouhse, Tingting Ju, Yi Fan, Camila S. Marcolla, Robert Pieper, Ryan K. Brook, Benjamin P. Willing

## Abstract

The microbiome of wild animals is believed to be co-evolved with host species, which may play an important role in host physiology. It has been hypothesized that the rigorous hygienic practice in combination with antibiotics and diets with simplified formulas used in the modern swine industry may negatively affect the establishment and development of the gut microbiome. In this study, we evaluated the fecal microbiome of 90 domestic pigs sampled from 9 farms in Canada and 39 wild pigs sampled from three different locations on two continents (North America and Europe) using 16S rRNA gene amplicon sequencing. Surprisingly, the gut microbiome in domestic pigs exhibited higher alpha-diversity indices than wild pigs (*P*<0.0001). The wild pig microbiome showed a lower Firmicutes to Bacteroidetes ratio and a higher presence of bacterial phyla Elusimicrobiota, Verrucomicrobiota, Cyanobacteria, and Fibrobacterota compared to their domestic counterparts. At the genus level, wild pig microbiome had enriched genera that were known for fibre degradation and short-chained fatty acids production. Interestingly, the phylum Fusobacteriota was only observed in domestic pigs. We identified 31 ASVs that were commonly found in the pig gut microbiome regardless of host sources, which could be recognized as members of the core gut microbiome. Interestingly, we found a few ASVs missing in domestic pigs that were prevalent in wild ones, whereas domestic pigs harbored 59 ASVs that were completely absent in wild pigs. The present study sheds light on the impact of domestication on the pig gut microbiome, including the gain of new genera.

**Importance:** The microbiome of pigs plays a crucial role in shaping host physiology and health. This study looked to identify if domestication and current rearing practices have resulted in a loss of co-evolved bacterial species by comparing the microbiome of wild boar and conventionally raised pigs. It represents a comparison of domestic and wild pigs with the largest sample sizes, and is the first to examine wild boars from multiple sites and continents. We were able to identify core microbiome members that were shared between wild and domestic populations, and counter to expectation, few microbes were identified to be lost from wild boar. Nevertheless, the microbiome of wild boars was distinct from domestic pigs, with notably lower abundance of important pathogenic genera. The differences in microbial composition may identify an opportunity to shift the microbial community of domestic pigs towards that of wild boar with the intent to reduce pathogen load.

## Background

Approximately ten million years ago, domestic pigs (*Sus domesticus*) underwent a divergence from their wild ancestors (*Sus scrofa*) in Eurasia (1, 2). In the late 1980s, domestic wild boar was introduced to Canada as a livestock diversification initiative (3). Subsequently, a subset of these animals either escaped or were deliberately released (4). These free-ranging wild pigs have now become established in the wild and have spread rapidly in Western Canada. While returning to the wild may exhibit certain ancestral traits (5, 6), it is not merely a straightforward reversal of domestication (7). This re-wilding of domesticated wild boars has introduced selective pressures, favoring the survival and reproduction of larger individuals to optimize overall fitness for colder weather, consistent with Bergmann’s rule in ecological theory (8). In addition to physical traits, the selection of processes applied during domestication has resulted in significant alterations in the gut microbiota composition (9, 10). Such changes in the gut microbial composition in domestic pigs have been linked to factors such as diets, antibiotic usage, reduced exposure to natural environments, and practices associated with intensive farming (11–13). Notably, wild pigs engage in extensive rooting behavior, which involves ripping up the soil to consume roots and insect larvae, resulting in a distinct soil-influenced gut microbial community structure (14).

It has been well recognized that the gut microbiome can significantly impact host health, metabolism, and immune function, playing a crucial role in shaping the symbiotic and co-evolutionary dynamics with the host organism (15). The profound interdependence between the host and its microbiota indicated a dynamic and reciprocal process of beneficial adaptation and evolution. Recent studies have indicated that the gut microbiota from wild mice can out-compete lab mouse microbiota and improve host resilience to infectious diseases (16, 17). Similarly, wild pigs show robust environmental adaptability and resistance to diseases compared with domestic pigs (10, 18–23), suggesting a potential link between the gut microbiome and observed differences in host phenotypes.

In addition to recent studies that have explored the gut microbiota in pigs focusing on agricultural traits and applications (24–28), a few studies explored the gut microbial compositions in wild pigs from different regions of the world (21, 22, 29). For example, comparative analyses of the gut microbial composition among wild boars, feral, and domestic pigs demonstrated differences in microbial profiles between juveniles and adults in Italy regardless of the population (30). Fungal communities in the pig gut across a production cycle have also been investigated, drawing distinctions between feral and farm pigs (31). Despite these efforts, there is still limited information about the microbial signature composition of domestic and wild pigs. The current study compared the gut microbiome between domestic and wild pigs from different sites and habitats across 2 continents. We hypothesized that the husbandry practices of pigs in the domestic environment have different microbiome signatures compared with wild pigs, and such differences may lead to a loss of beneficial co-evolved gut microbes that could potentially be used for future efforts to improve disease resilience.

## Methods

### Animals and sample collections

We sampled 9 farms in Western Canada, including 7 from Alberta and 2 from British Columbia, to characterize the microbial communities of domestic pigs. All farms used disinfectants for barn sanitation; only one used prophylactic antibiotics (Excede). The domestic pig populations in the study originated from five different commercial breeders and genetic companies. Herd sizes varied considerably, ranging from 50 to 2,800 sows among the farms. Additionally, the weaning age of piglets across the farms ranged from 20 to 30 days. Among the farms, the nursery (post-weaning; 4-8 weeks of age) and grower (8-12 weeks of age) mortality rates range from 1% to 3.8% and 1.7% to 5.6%, respectively. All domestic pigs were housed on slatted floors, and all but one farm fed a creep diet. Faecal swabs were collected from 20-week-old pigs and sows in their second parity (n = 5 per group, totaling N = 90 samples). For wild pig populations, we collected samples from Canadian and German wild pigs. For Canadian wild pig populations, 34 wild pigs of reproductive age were captured in Melfort (n = 24) and Moose Mountain (n = 10), Saskatchewan, with a net gun after being located by helicopter. Captured pigs were subsequently euthanized via a captive bolt, and distal colon digesta was collected and snap-frozen for downstream analyses. For the German wild pig population, reproductive-age wild boars (n = 5) at a driven hunt approximately 80 km north of Berlin were field-dressed at a central point, and colon digesta samples were taken aseptically from eviscerated organs (32).

### Microbial DNA extraction and 16S rRNA gene amplicon sequencing

Microbial DNA extraction was carried out using the DNeasy PowerSoil kit (Qiagen®, Valencia, CA) following the manufacturers’ instructions with an addition of a bead-beating step on a FasPrep-24^TM^ (MP Biomedicals, OH, USA) homogenizer (5 m/s for 1 minute). The concentration of extracted DNA was determined using a Quant-iT™ PicoGreen® dsDNA Assay Kit (Thermo Fisher Scientific, Waltham, MA, USA). Amplicon libraries were generated using the Illumina^®^ 16S metagenomic sequencing library preparation protocol, targeting the V3-V4 region of the 16S rRNA gene, using PCR primers 341F (5’-TCGTCGGCAGCGTCAGATGTGTATAAGAGACAGCCTACGGGNGGCWGCAG-3’) and 785R (5’-GTCTCGTGGGCTCGGAGATGTATAAGAGACAGGACTACHVGGGTATCTAATCC-3’). A 2 x 300 bp paired-end sequencing was performed on an Illumina MiSeq platform (Illumina, Inc., Sandiego, CA).

Sequencing data was processed using Quantitative Insights Into Microbial Ecology 2 (QIIME2; v2020.6) (33). The FastQC with default parameters was used to assess the quality of sequences, and low-quality reads were filtered by a quality score of 20 as a cutoff value. The Divisive Amplicon Denoising Algorithm 2 (DADA2) plugin was used to denoise and generate feature tables (34) with truncation lengths of 270 and 210 bp for forward and reverse reads, respectively. Amplicon sequence variants (ASVs) were aligned with mafft program (35) and subsequently used to generate a phylogenetic tree with FastTree 2 (36). Taxonomic assignment for the ASVs was made using the QIIME2 q2-feature-classifier (37) and the classify-sklearn I Bayes taxonomy classifier (38) using the pre-trained nearly complete 16S rRNA sequences from the SILVA database (version 138) (39).

### Data visualization and statistical analysis

All statistical analyses were conducted with RStudio (v3.5.2) and GraphPad Prism (v9.5.1, GraphPad Software Inc, San Diego, USA). Data are shown as mean ± standard error of the mean. The phyloseq package (40) in R was used to analyze microbial community structure and alpha diversity indices. Rarefaction was based on the sample with the lowest sequence depth (6,000 read count) for diversity measurements. Shannon, Observed features, and Pielou evenness indices were analyzed with Mann-Whitney U or Kruskal-Wallis tests, followed by post-hoc Dunn tests for significance analyses in GraphPad Prism. Differences in microbial community compositions were analyzed by PERMANOVA based on Bray-Curtis dissimilarities and visualized by principal-coordinate analysis (PCoA). Differentially abundant taxa at the genus level were calculated with a Wald parametric test with Benjamini-Hochberg adjustment in the DESeq2 package in R (41). Statistical significance was set at a *P*-value less or equal to 0.05. ASVs derived from domestic pigs, wild pigs, or shared between the two were summarized. An ASV identified in domestic pigs was classified as ‘Domestic Origin’ if the ASV was present in more than 50% of the domestic pig samples but absent in wild pigs. Conversely, an ASV was classified as ‘Wild Origin’ if the ASV was not identified in conventional pigs but was found in more than 50% of wild pigs. An ASV was classified as ‘Shared’ when identified in more than 50% of samples in both groups. The ggplot2 package in R (v3.5.2) was utilized to visualize the presence of ASVs.

## Result

To explore the differences in the gut microbial composition between wild and domestic pigs, a total of 4,742,456 raw reads were generated, averaging 36,763 ± 2,290 (SEM) reads per sample. Following the removal of samples with low reads (n = 8) and quality control, the average read count was 37,093 ± 2,345 (SEM) reads per sample. The rarefaction curves were evaluated based on Shannon and Pielou evenness, and observed features indices indicated adequate reads for the analyses as saturation curves reached a plateau.

### The gut microbial community structure in wild pigs is different from domestic pigs

Domestic pigs exhibited greater species richness compared to wild pigs, as evidenced by observed features (Fig. 1A). Domestic pigs also showed a higher Shannon diversity index compared to their wild counterparts (Fig. 1B), while Pielou indices, which only measures the evenness of microbial community distribution, was not different (Fig. 1C). We found alpha diversity based on Shannon and observed features significantly different among the farms, while no differences were observed in Pielou indices (Supplementary Figure 1A-C). Geographical locations did not impact Shannon and observed features indices in wild pig populations (Supplementary Figure 2A-B). However, evenness based on Pielou indices varied among wild pig populations across different locations (Supplementary Figure 2C). In domestic pigs, there were no differences in the two age groups for alpha diversity indices (Supplementary Figure 3A-C).

**Figure 1:**
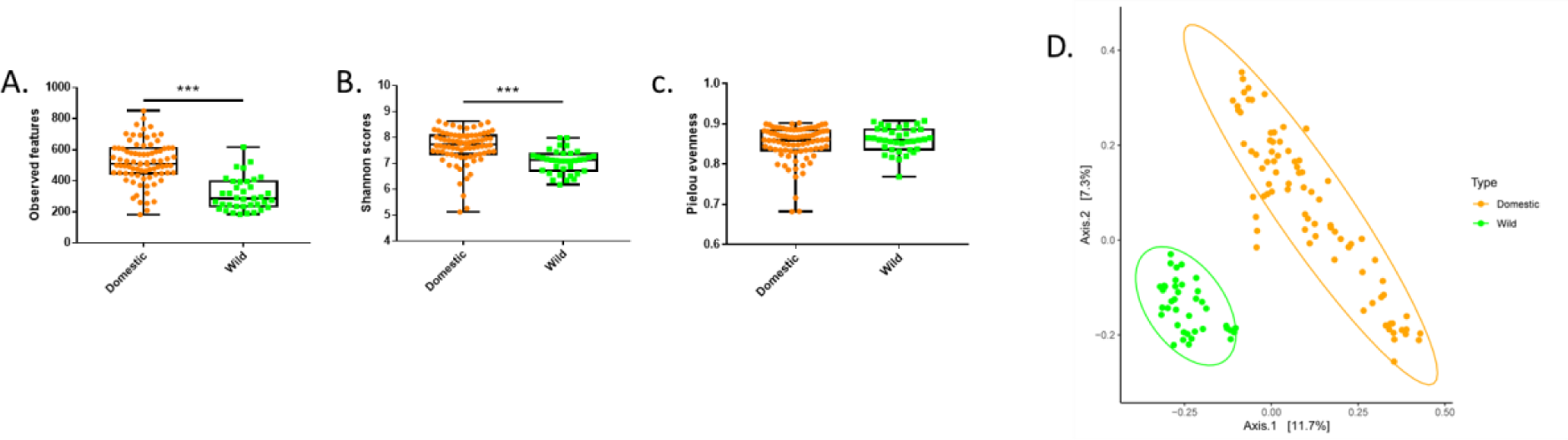
Comparisons of fecal microbial community structure and α-diversity indices between domestic (n = 82) and wild pigs (n = 39). (A-C) Domestic pigs have higher species richness (*P* < 0.0001) and diversity (*P* < 0.0001) but no differences in evenness (*P* = 0.52) compared to their wild counterparts. (B) The microbial community structure was different as measured based on Bray–Curtis dissimilarity matrix (Adonis *P* = 0.001, R^2^ = 0.096; Betadispersion *P* = 0.219).

The difference in microbial community structures between domestic and wild pigs was analyzed based on Bray-Curtis dissimilarities (Fig. 1D). A homogeneous dispersion in both groups was observed by beta-dispersion analysis (*P* = 0.219). There was a clear separation between the gut microbial communities between wild and domestic pigs (Adonis *P* = 0.001, R^2^ = 0.096). In domestic pigs, we observed a significant farm (Adonis *P* = 0.001, R^2^ = 0.203; beta-dispersion *P* = 0.96) and age (Adonis *P* = 0.001, R^2^ = 0.089; beta-dispersion *P* = 0.90) effect in microbial community structures (Supplementary Figure 3-4). In wild pigs, microbial communities varied (Adonis *P* = 0.001, R^2^ = 0.257; beta-dispersion *P* = 0.31) by location (Supplementary Figure 5), however, the German wild boar grouped closely with the Canadian wild boar relative to the variations in domestic population.

### Microbial taxonomic composition of domestic and wild pigs

The taxonomic profiling of the gut microbiota in wild and domestic pigs was analyzed, which identified 13 bacterial phyla in wild and domestic pigs with a relative abundance greater than 0.5% (Fig. 2A). In domestic pigs, Firmicutes (44.5%) was the most predominant phylum, followed by Bacteroidetes (40.9%), Spirochaetota (4.5%), Proteobacteria (3.6%), Campilobacterota (1.7%), and Actinobacteriota (1.2%). While in wild pigs, Bacteroidetes (52.9%) was the predominant phylum, succeeded by Firmicutes (36.3%), Spirochaetota (3.1%), Verrucomicrobiota (2.6%), Proteobacteria (1.9%), and Fibrobacterota (0.8%). Interestingly, the phylum Fusobacteriota was exclusively observed in domestic pigs. Domestic pigs’ gut microbiota was enriched (Mann-Whitney U test, FDR <0.05) with the phyla Campilobacterota, Actinobacteriota, Spirochaetota, and Proteobacteria, whereas the wild pigs’ gut microbiota was enriched with Elusimicrobiota, Verrucomicrobiota, Fibrobacterota, and Cyanobacteria (Fig. 2A).

**Figure 2:**
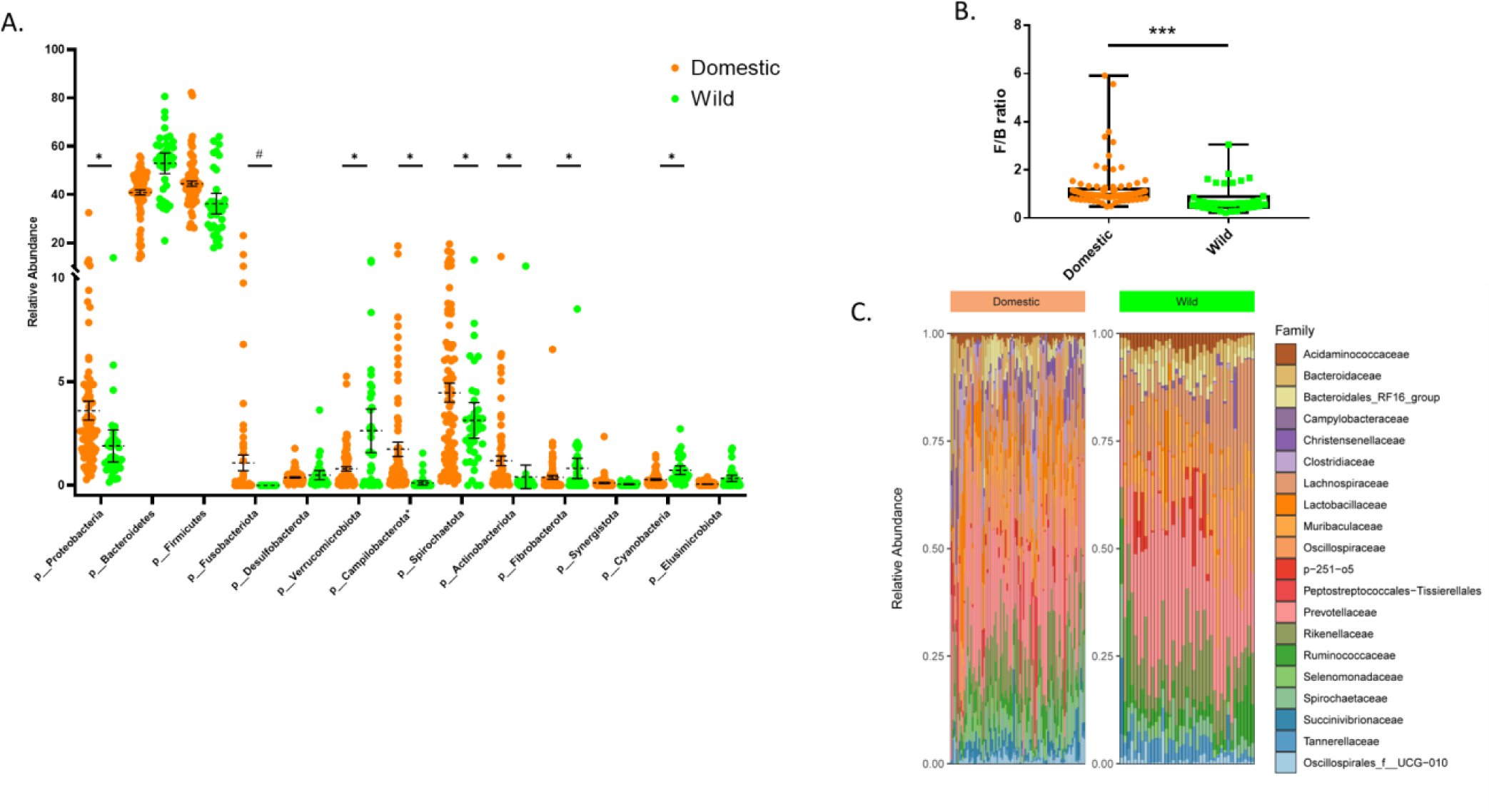
Taxonomical profiling based on relative abundance revealed that the microbial composition of domestic and wild pigs differed. (A) Scatter plot (mean ± SEM) showing the composition of top 13 bacterial phyla observed between domestic and wild pigs (Asterisk (*) indicates significant differences between groups; while pound (#) indicates the phyla only present in the domestic group). (B) Box and whiskers plot (Minimum to Maximum) illustrating that wild pigs exhibit a lower Firmicutes to Bacteroidetes ratio. (C) Top 20 bacterial families observed in domestic and wild pigs.

At the family level, 61 families were identified in both domestic and wild pig samples combined (Supplementary Table 1), considering > 0.1% of the relative abundance of the gut microbiota in each group. Both wild and domestic pigs’ gut microbiota shared the same pattern of the top 5 families (Fig. 2C), which were *Prevotellaceae* (domestic 19.4% vs. wild 23.1%), *Lachnospiraceae* (domestic 6.9% vs. wild 12.3%), *Oscillospiraceae* (domestic 6.7% vs. wild 9.2%), *Rikenellaceae* (domestic 6.7% vs. wild 8.18%), and *Muribaculaceae* (domestic 4.8% vs. wild 5.5%). Among the 61 families, 9 families including *Peptostreptococcales-tissierellales*, *Porphyromonadaceae*, *Fusobacteriaceae*, *Aerococcaceae*, *Moraxellaceae*, *Planococcaceae*, *Corynebacteriaceae*, *Mycoplasmataceae*, and *Carnobacteriaceae* were only observed in domestic pigs (Supplementary Table 1).

At the genus level, the most abundant genera of the fecal microbiota of domestic pigs were *Prevotella*, *Lactobacillus*, *Clostridium_sensu_stricto_1*, *Christensenellaceae_R-7_group*, and *Fusobacterium* (Fig.3). In the wild pig population, *Prevotellaceae_NK3B31_group*, an unknown genus of *Lachnospiraceae*, *Alloprevotella*, *Rikenellaceae_RC9*, *Parabacteroides*, and *Ruminococcus* were ranked as the most abundant bacterial genera.

**Figure 3:**
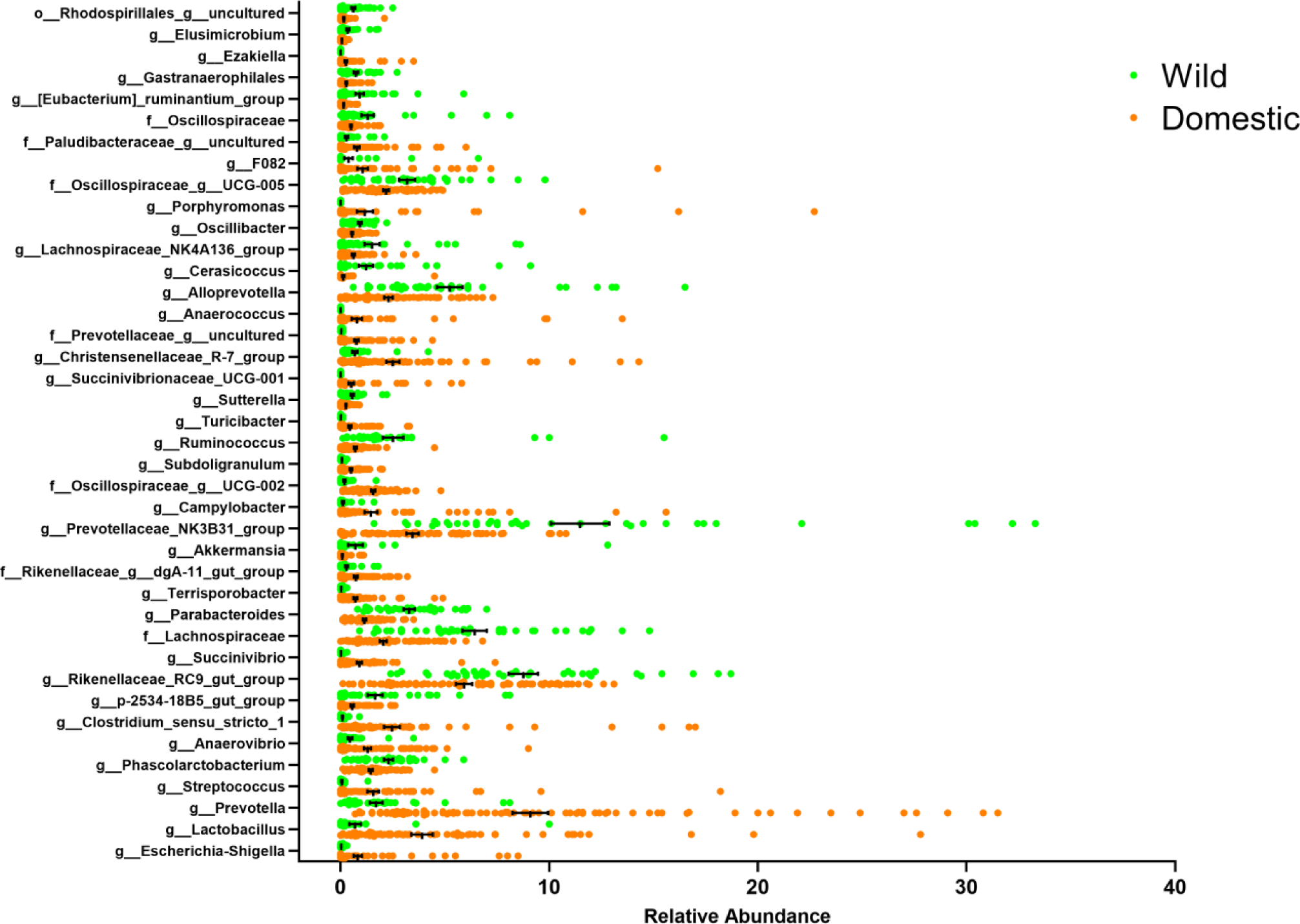
The relative abundance of bacterial genera that were differentially abundant between domestic and wild pigs was determined using the Mann-Whitney test followed by an FDR adjustment. To increase the accuracy of identifying the differentially abundant taxa, features that exhibited more than 2 log LDA differences and were significant after an FDR adjustment of 0.05 in LEfSe and DESeq2 were included in the graphs.

### Core microbiome of domestic and wild pigs

ASVs present in over 50% of samples of each group were selected to identify core members exclusively in domestic and wild pigs. Among the selected ASVs, 59 were exclusively present in domestic pigs (Fig. 4) with 29 ASVs attributed to Firmicutes, 20 ASVs attributed to Bacteroidota (14 were linked to the genus *Prevotella*), 4 belonging to Spirochaetota (the genus *Treponema*), and 2 belonging to Proteobacteria. Conversely, in wild pigs, only 5 ASVs were detected in over 50% of samples, which were absent in domestic pigs. Three of these ASVs belong to the Firmicutes (two were associated with *Oscillibacter* and one was characterized as *Staphylococcus*), and one ASV belong to Bacteroidota, as well as one was attributed to Desulfobacterota.

**Figure 4:**
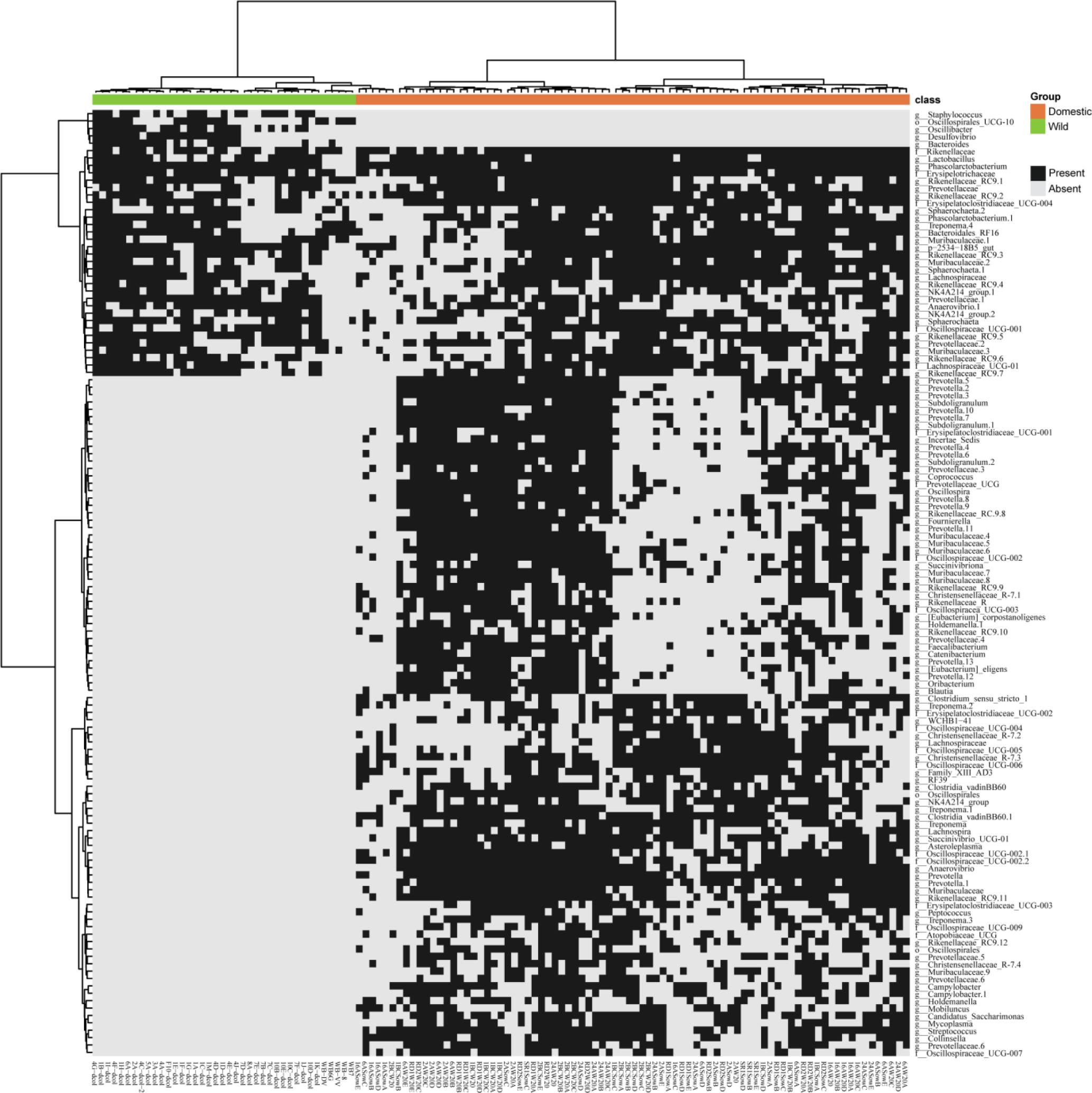
Heatmap representation of domestic and wild pigs’ shared and unique ASVs based on the presence or absence of each ASV. For the core microbiome, the frequency of each ASV was present in >50% of the samples of each group. The ASVs were agglomerated at the genus level (when this classification is available) and are specified on the left, and the sample IDs are on the bottom and groups on the top. The Ward hierarchical clustering algorithm based on Euclidean distances was used to cluster the ASVs and the samples. Black indicates the presence of the ASV, while gray indicates absence.

To determine core members across both domestic and wild pigs, 31 ASVs were identified and present in over 50% of the samples of each group. Among these, 16 ASVs belonged to Bacteroidota (*Rikenellaceae_RC9*, *Prevotella*, and *Muribaculaceae*). Additionally, 11 ASVs originated from the phylum Firmicutes, including members from *Oscillospiraceae*, *Lachnospiraceae*, *Phascolactebacterium*, *Lactobacillus*, *Erysipelatoclostridiaceae*, and *Anaerovibrio* (Fig. 4).

## Discussion

Gut microbial communities have shown a multifaceted and significant influence on host biology. They coevolve with their host and play a crucial role in the host’s ability to adapt to its environment and link to health and disease (17, 42). The gut microbiome of wild animals has been linked with improving host immune development and resilience to diseases compared to domesticated counterparts, as evidenced in several studies involving different animal models (16, 43–45). Several studies have been conducted to explore the differences between the gut microbiota of wild and domestic pigs (25, 30, 46, 47). However, these studies were limited by sample size, number of sites, and geographical locations. The present study investigated the fecal microbiota of domestic pigs from 9 different farms in Canada and wild pigs from three habitats across two continents (North America and Europe). We report that domestic pigs have higher microbial richness and diversity, as well as altered microbial community structures than their wild counterparts. In addition, contrary to our original hypothesis, domestic pigs exhibited a higher presence of unique ASVs than wild pigs.

In line with previous studies, the current study demonstrated differences in the gut microbial compositions between domestic and wild pigs (22, 30, 46, 48). It has been documented that rearing conditions and diets are the primary drivers of gut microbial composition. Domestic pigs are reared indoors with rigorous hygienic practices and different diets (24, 49–51). Therefore, the differences in gut microbiota structures between domestic and wild pigs are expected. Interestingly, we observed lower alpha diversity indices representing richness and diversity in wild pigs than that in domestic pigs. Consistent with our findings, a recent study found a trend of lower bacterial richness in adult wild pigs compared to domestic pigs (30). Likewise, many other studies across different species observed either no difference or higher microbial richness in captive or domestic animals compared to the wild ones (9, 52–54). The distinct differences observed between domestic and wild pig microbial community structure and composition are likely influenced by several factors, including host specialization, environmental pressures, and the process of natural selection (55). Furthermore, the dietary habits of wild pigs, which often consist of complex non-digestible carbohydrates sourced from plants, could serve as another significant driver behind the observed decrease in alpha diversity and the development of specific microbial communities within wild pig populations. Notably, previous research has demonstrated that the supplementation of swine feed with resistant starch reduced microbial diversity while concurrently promoting the abundance of beneficial bacteria (56).

Our study found that Firmicutes and Bacteroidetes dominated the gut microbiota of both domestic and wild pigs with different ratios. The higher Bacteroidetes in wild pigs was consistent with previous studies exploring the gut microbiota of healthy wild boars (27), which has also been observed in extensively raised chickens (57) and in humans living in rural areas (58, 59). It has been shown that Bacteroidetes are capable of breaking down various types of complex non-digestible carbohydrates in the host gastrointestinal tract, while Firmicutes are well-suited to the cross-feeding process in the gut (60). Lower F/B ratios in pigs have previously been linked to a regular diet without antibiotics compared to diets with antibiotics and essential oils, which was associated with lower body weight and average daily gain (54). Another study using a high-resolution metagenomics approach showed that increased fat deposition in pigs was associated with a higher F/B ratio along with other microbial signatures (61). Likewise, the microbiome of obese mice demonstrates a higher F/B ratio and an increased capacity to extract energy from the diet compared to lean individuals (62). These findings indicated a potential association of altered F/B ratio with caloric restriction. Indeed, according to a study in brown bears, the shift from an active lifestyle in summer to hibernation in winter was linked to a rise in Bacteroidetes and a decline in Firmicutes (63). Together, these results infer that the alteration in the F/B ratio of domestic pigs might be attributed to diets and the fact that wild pigs lead a more energetically demanding lifestyle.

Interestingly, domestic pigs exhibited a higher abundance of Campilobacterota, Actinobacteriota, Spirochaetota, Proteobacteria, and the exclusive presence of Fusobacteriota phyla. At the genus level, the gut microbiota of domestic pigs was also enriched with potential pathogenic genera such as *Fusobacterium*, *Porphyromonas*, *Campylobacter*, *Streptococcus*, *Treponema*, and *Escherichia-Shigella.* These genera include numerous potential pathogens, which were not only of concern for pig health but also a risk for human food-borne disease. On the other hand, a higher presence of Elusimicrobiota, Verrucomicrobiota, Fibrobacterota, and Cyanobacteria in wild pig gut indicates an influence of the environment and a greater capability for fiber degradation (20, 64–67). Genus-level differential analyses revealed that wild pig microbiomes were enriched with beneficial bacterial genera capable of fiber degradation and SCFA production. Notable enriched fiber fermenting taxa included members from the families *Preveotellaceae*, *Lachnospiraceae*, *Osillospiraceae*, and *Muribaculaceae*, as well as the genera *Alloprevotella*, *Rikenellaceae_RC9*, *Ruminococcus*, *Bacteroides*, *Fibrobacter*, *Elusimicrobium,* and *Fecalibacterium* (68). One potential explanation for the low abundance of beneficial bacterial genera in domestic pigs could result from rigorous hygienic protocols and confinement within indoor rearing environments. These practices may unintentionally impede the natural colonization of beneficial bacteria. Conversely, when pigs are reared outdoors, they root in the soil, which has been shown to enhance the colonization of beneficial microbes (14, 69). Indeed, prior research has demonstrated that indoor rearing conditions and stringent hygiene measures tend to elevate the prevalence of certain potentially harmful bacterial genera such as *Campylobacter*, *Streptococcus*, *Fusobacteria*, *Treponema*, and *Escherichia/Shigella* in domestic pigs (70, 71). It has also been observed in poultry production that increased levels of sanitation were associated with increased *Campylobacter* loads (72). Conversely, outdoor rearing environments have been associated with a higher abundance of beneficial bacteria like *Faecalibacterium*, *Elusimicrobium*, and *Ruminococcus,* which have been reported to regulate gut health and energy metabolism through the production of SCFAs (14, 49, 73). Interestingly, the majority of the differentially abundant genera in domestic pigs are commonly found in the human gut and are integral members of the human core microbiome (74). This shared microbial composition suggests a potential intermingling or encroachment of the human microbiome during the domestication process (9). As a result, the heightened presence of these phyla in farm pigs could reflect a convergence of microbial communities between humans and domesticated animals, influencing the composition of the gut microbiota in pigs reared in farm settings.

Consistent with other studies, we observed the dominance of *Prevotella* in domestic and wild pigs (14, 46, 47, 51). *Prevotella*, a core member of the pig large intestine microbiota (75), is actively linked with carbohydrate metabolism and has shown positive correlations with enhanced feed efficiency, weight gain, and reduced incidence of diarrhea (76). Our study identified distinct *Prevotella* predominance between domestic and wild pigs, with wild pigs exhibiting a higher abundance of the genus *Prevotellaceae_NK3B31_group*, whereas domestic pigs showed a prevalence of *Prevotella* and an uncultured member of the *Prevotellaceae* family. These findings parallel those of prior studies, indicating a consistent pattern of differing *Prevotella* dominance in wild and domestic pig populations (30). Moreover, earlier research has underscored the fluctuations in different *Prevotella* compositions in the pig gut before and after weaning, suggesting dietary modifications as a potential driver of this phenomenon (77, 78). Similarly, recent investigations in human research have suggested that the selection of distinct *Prevotella* species in western and non-western populations was primarily influenced by diets and lifestyle (79). Therefore, the observed variations in the predominance of different *Prevotella* in wild and domestic pigs likely stem from differences in dietary composition and environmental conditions.

We found that 31 ASVs were conserved between domestic and wild pigs, most of which were members of the phyla Bacteroidetes and Firmicutes. Among them, 29 ASVs were consistent with the core microbiome members reported in a previous study (25); the only exceptions were *Phascolarctoebacterium* and *Erysipelatoclostridiaceae*. This observation implies that gut bacteria are symbionts that have maintained a stable association with their hosts, even after the wild and domesticated populations split. Five ASVs characterized as *Oscillibacter*, *Staphylococcus*, *Desulfovibrio*, and *Bacteroides* were exclusively found in wild pigs and might be members of ancestral microbiomes that were lost during domestication. Meanwhile, 59 ASVs exclusively observed in domestic pigs were missing in wild pigs. Acquiring these unique ASVs indicates the plasticity and adaptability of the swine gut microbiome in response to environment, rearing conditions, and diet (53, 80, 81).

The presence of several limitations in the current study underscores the need for careful consideration of the results. First, we compared the fecal swab microbiota of domestic pigs with the distal colon microbiota of wild pigs, although the differences can be negligible (82). Second, the microbial taxonomic assignment of the current study was based on 16S rRNA gene amplicon sequences, which limits the resolution to classify taxa at the species level and does not provide functional profiling. Therefore, a high-resolution metagenomic approach to resolve taxonomic assignment and functional capacity is warranted.

This study showed significant differences in the gut microbiome between domestic and wild pig populations. To a great extent, these variations were characterized by increased bacterial diversity and altered microbial community compositions in domestic pigs. Additionally, we found that the gut microbiomes of wild pigs, adapted to the lifestyle in the wild, were enriched with phyla known to be involved in caloric restriction, the breakdown of complex fiber, and the production of SCFAs. Despite the theory that low dispersal and distinct selective environments with antibiotics and low fiber intake can reduce successful colonization rates and may lead to bacterial lineage extinction (83, 84), we observed a small number of bacterial members found in the wild pig gut missing in domestic pigs. Our study provides insight into gut microbial compositions impacted by the pig domestication process, which can be used in future microbial modulation to improve health and disease resilience. Further studies could help verify these microbial signatures and examine the impact on metabolic and immune function between wild and domestic pig populations.

## Data Availability

Raw reads from the16S rRNA gene amplicon sequencing was deposited to the National Center for Biotechnology Information Sequence Read Archive and is available under BioProject accession number PRJNA1091615.

## Authors’ contributions

B.P.W. conceived and designed the study and secured the funding. J.M.F. oversaw the data and sample collection process, with help from R.K.B. and R.P. R.R. and J.M.F. performed the laboratory work. R.R. and T.J. performed bioinformatic and statistical analyses. R.R. wrote and revised the main manuscript with input from all authors. All authors read and approved the final manuscript.

## Acknowledgments

The Alberta Livestock and Meat Agency (res0030386) and a Natural Sciences and Engineering Research Council of Canada Discovery grant (RGPIN-2019–06336) provided funding for this research. B.P.W. received support from the Canada Research Chair Program, while R.R. was supported by an Alberta Graduate Excellence Scholarship and Frank Aherne Graduate Scholarship in Swine Research. We extend our gratitude to all the staff of participating farms and the Swine Research and Technology Centre of the University of Alberta for their invaluable help and support. Additionally, we acknowledge the University of Saskatchewan and the United States Animal and Plant Health Inspection Service National Feral Swine Damage Management Program for funding the wild pig captures.

---

**Supplementary Table 1:** The top 61 bacterial families (cut-off, 0.1% relative abundance in domestic and wild pigs) were observed in both groups. Families that were only present in the domestic pigs were labeled with the “#” sign.

| Family | Mean Relative Abundance (%) |  |
| --- | --- | --- |
|  | Domestic pigs | Wild Pigs |
| f__Prevotellaceae | 19.366 | 23.059 |
| f__Lachnospiraceae | 6.937 | 12.321 |
| f__Oscillospiraceae | 6.736 | 8.184 |
| f__Rikenellaceae | 6.690 | 9.231 |
| f__Muribaculaceae | 4.811 | 5.524 |
| f__Spirochaetaceae | 4.466 | 3.128 |
| f__Lactobacillaceae | 3.917 | 0.693 |
| f__Ruminococcaceae | 2.813 | 4.330 |
| f__Christensenellaceae | 2.505 | 0.684 |
| f__Clostridiaceae | 2.484 | 0.102 |
| f__Bacteroidaceae | 2.151 | 3.008 |
| f__Selenomonadaceae | 1.963 | 0.515 |
| f__Bacteroidales_RF16_group | 1.812 | 2.684 |
| f__Peptostreptococcales-Tissierellales# | 1.715 | 0.000 |
| f__Streptococcaceae | 1.563 | 0.052 |
| f__Acidaminococcaceae | 1.533 | 2.324 |
| f__Succinivibrionaceae | 1.486 | 0.607 |
| f__Campylobacteraceae | 1.448 | 0.108 |
| o__Oscillospirales_f__UCG-010 | 1.354 | 1.173 |
| f__Erysipelotrichaceae | 1.346 | 0.579 |
| f__Erysipelatoclostridiaceae | 1.202 | 0.680 |
| f__Porphyromonadaceae# | 1.163 | 0.000 |
| f__Peptostreptococcaceae | 1.160 | 0.051 |
| f__Clostridia_vadinBB60_group | 1.144 | 0.849 |
| f__Tannerellaceae | 1.140 | 3.281 |
| f__Veillonellaceae | 1.098 | 0.620 |
| f__Fusobacteriaceae# | 1.080 | 0.000 |
| f__F082 | 1.046 | 0.383 |
| f__p-251-o5 | 0.996 | 3.020 |
| f__[Eubacterium]_coprostanoligenes_group | 0.965 | 0.706 |
| f__Anaerovoracaceae | 0.942 | 0.538 |
| f__Enterobacteriaceae | 0.818 | 0.034 |
| f__Paludibacteraceae | 0.772 | 0.278 |
| f__Clostridia_UCG-014 | 0.713 | 0.405 |
| f__Actinomycetaceae | 0.562 | 0.002 |
| f__p-2534-18B5_gut_group | 0.556 | 1.649 |
| f__WCHB1-41 | 0.425 | 0.551 |
| f__Fibrobacteraceae | 0.381 | 0.815 |
| f__Aerococcaceae# | 0.319 | 0.000 |
| f__Moraxellaceae# | 0.314 | 0.000 |
| f__RF39 | 0.309 | 0.092 |
| f__Butyricicoccaceae | 0.283 | 0.329 |
| f__Gastranaerophilales | 0.270 | 0.725 |
| f__Helicobacteraceae | 0.262 | 0.010 |
| f__Desulfovibrionaceae | 0.259 | 0.358 |
| f__Sutterellaceae | 0.254 | 0.559 |
| f__Planococcaceae# | 0.251 | 0.000 |
| f__Corynebacteriaceae# | 0.211 | 0.000 |
| o__Bacteroidales | 0.184 | 0.779 |
| f__Monoglobaceae | 0.168 | 0.230 |
| f__Mycoplasmataceae# | 0.167 | 0.000 |
| o__Rhodospirillales;f__uncultured | 0.161 | 0.605 |
| f__Puniceicoccaceae | 0.144 | 1.292 |
| f__Izemoplasmatales | 0.142 | 0.028 |
| f__Pasteurellaceae | 0.141 | 0.044 |
| f__Acholeplasmataceae | 0.139 | 0.214 |
| f__Bifidobacteriaceae | 0.134 | 0.318 |
| f__Carnobacteriaceae# | 0.130 | 0.000 |
| f__Pseudomonadaceae | 0.124 | 0.003 |
| f__Bradymonadales | 0.117 | 0.130 |
| f__Synergistaceae | 0.113 | 0.053 |
| Other | 2.142 | 2.069 |

**Supplementary Figure 1:**
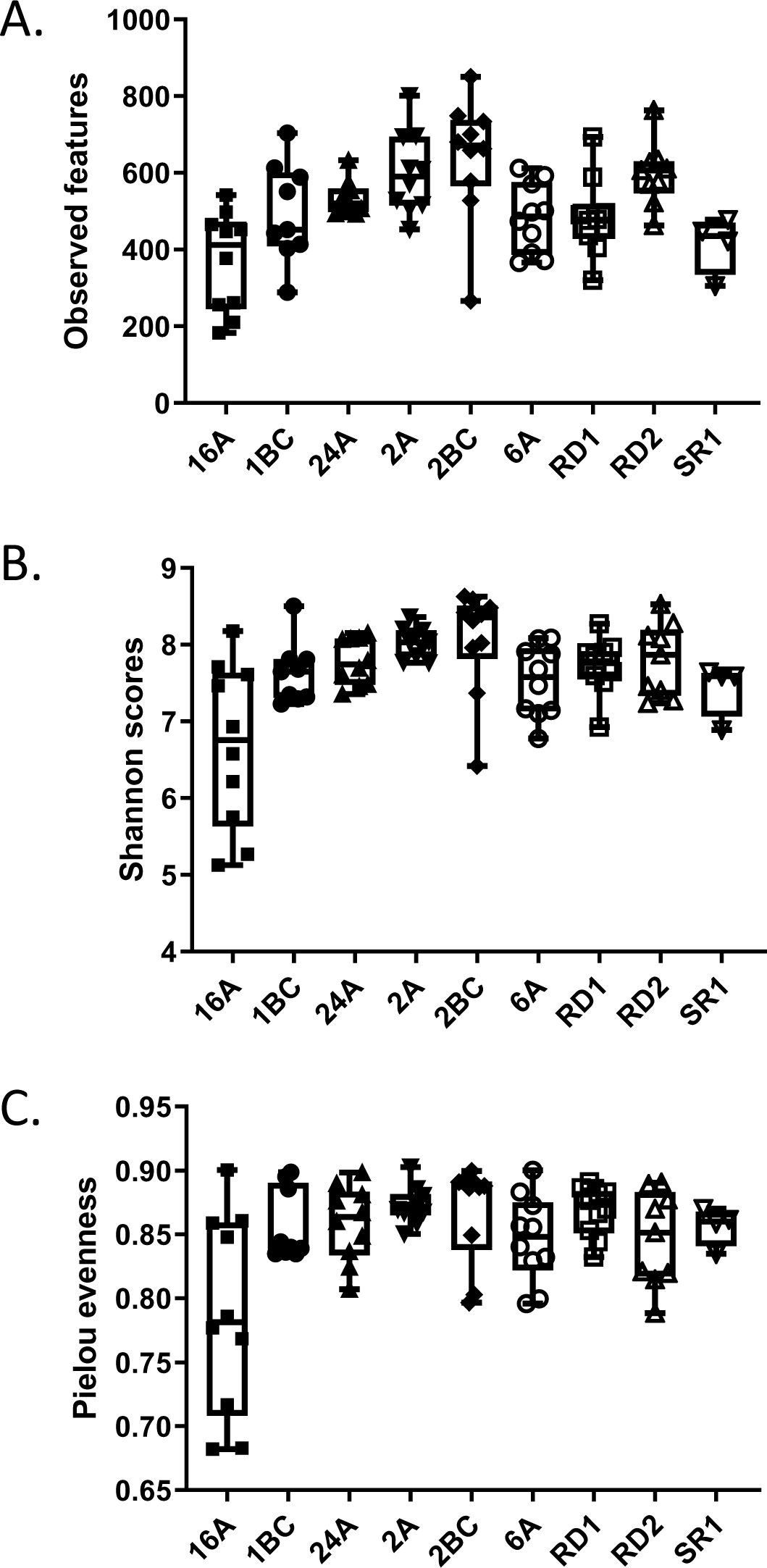
Comparisons of fecal microbial α-diversity indices among the 9 domestic farms using the Kruskal-Wallis test. (A) Based on the Observed features, the microbial richness analysis exhibited significant differences (*P* < 0.0001) across the farms. (B) Microbial diversity measured by Shannon index also showed significant variations (*P* = 0.002) among farms. (C) A trend in microbial evenness based on Pielou evenness scores among the domestic farms (*P* = 0.07). The farms in Alberta include 16A, 24A, 2A, 6A, RD1, RD2, and SR1, while those in British Columbia are 1BC and 2BC.

**Supplementary Figure 2:**
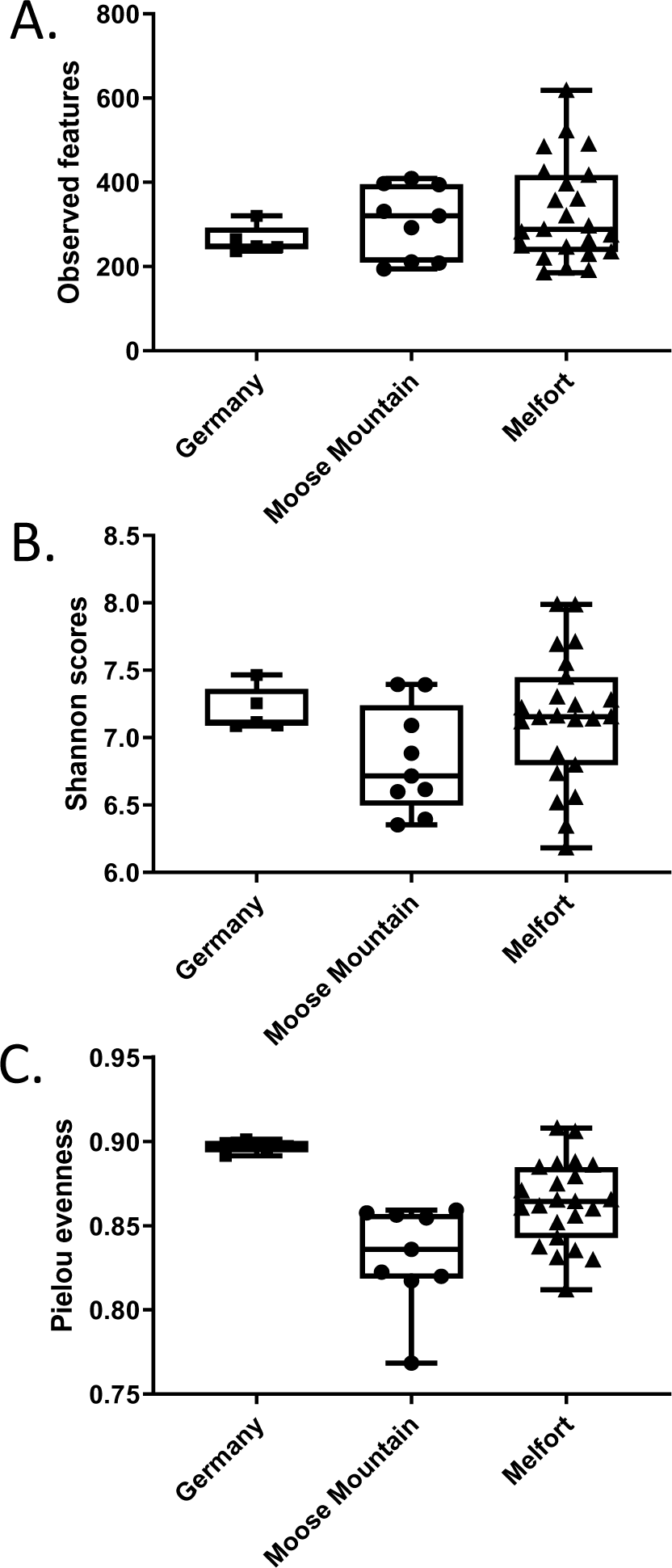
Comparisons of α-diversity indices of colonic microbial communities among the wild pigs from three locations using the Kruskal-Wallis test. No differences were observed for microbial richness (*P* = 0.61) based on the Observed features (A) and microbial diversity (*P* = 0.17) based on Shannon index (B). However, there were significant differences in the evenness (*P* = 0.0004) based on Pielou scores among the wild pigs’ originating locations (C).

**Supplementary Figure 3:**
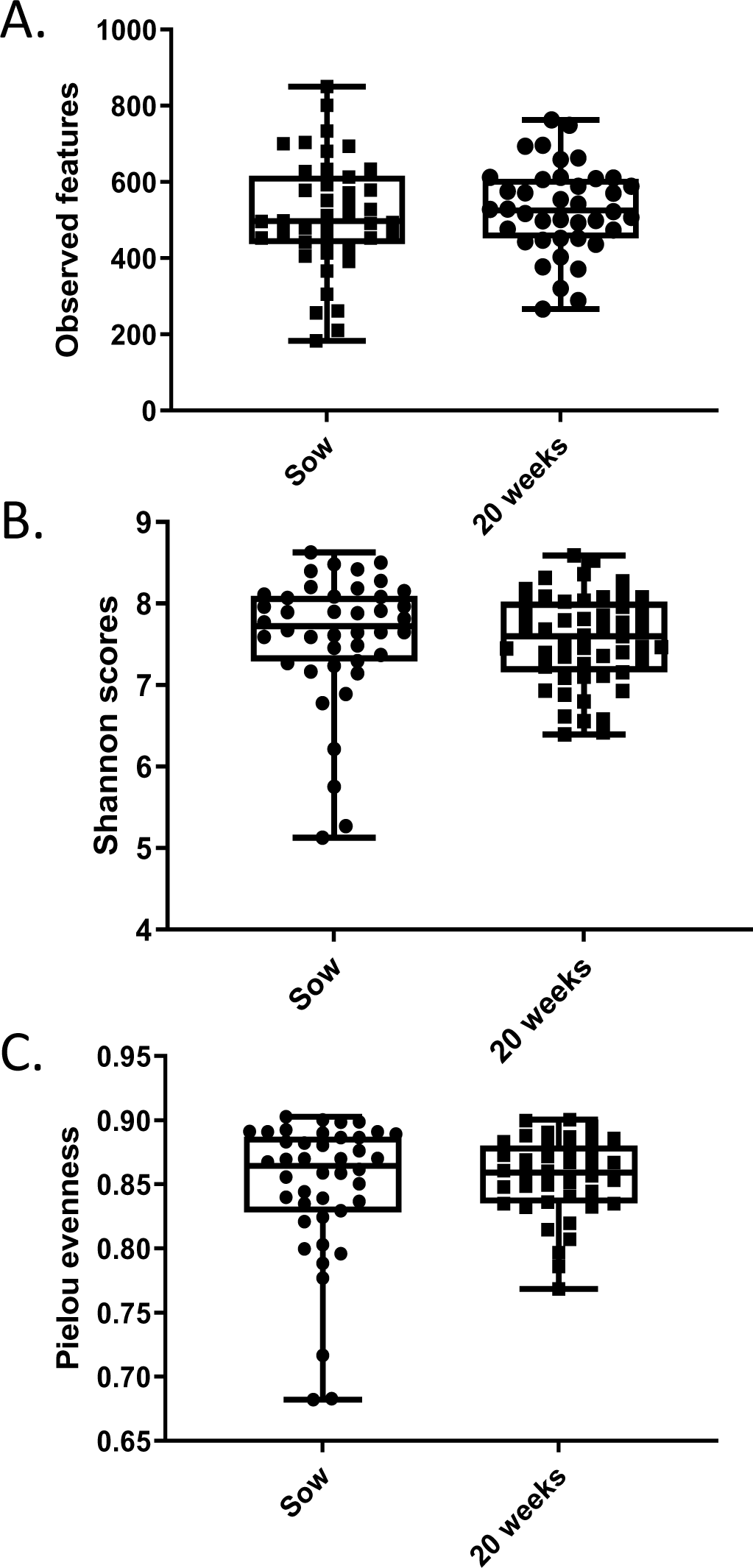
Comparisons of fecal microbial α-diversity indices between the Sow and 20-week groups of domestic pigs using the Mann-Whitney test. No differences were observed for microbial richness (*P* = 0.6) based on the Observed features (A), microbial diversity (*P* = 0.24) based on Shannon scores (B), or evenness (*P* = 0.76) based on Pielou evenness scores(C) between two age groups of the domestic pigs.

**Supplementary Figure 4:**
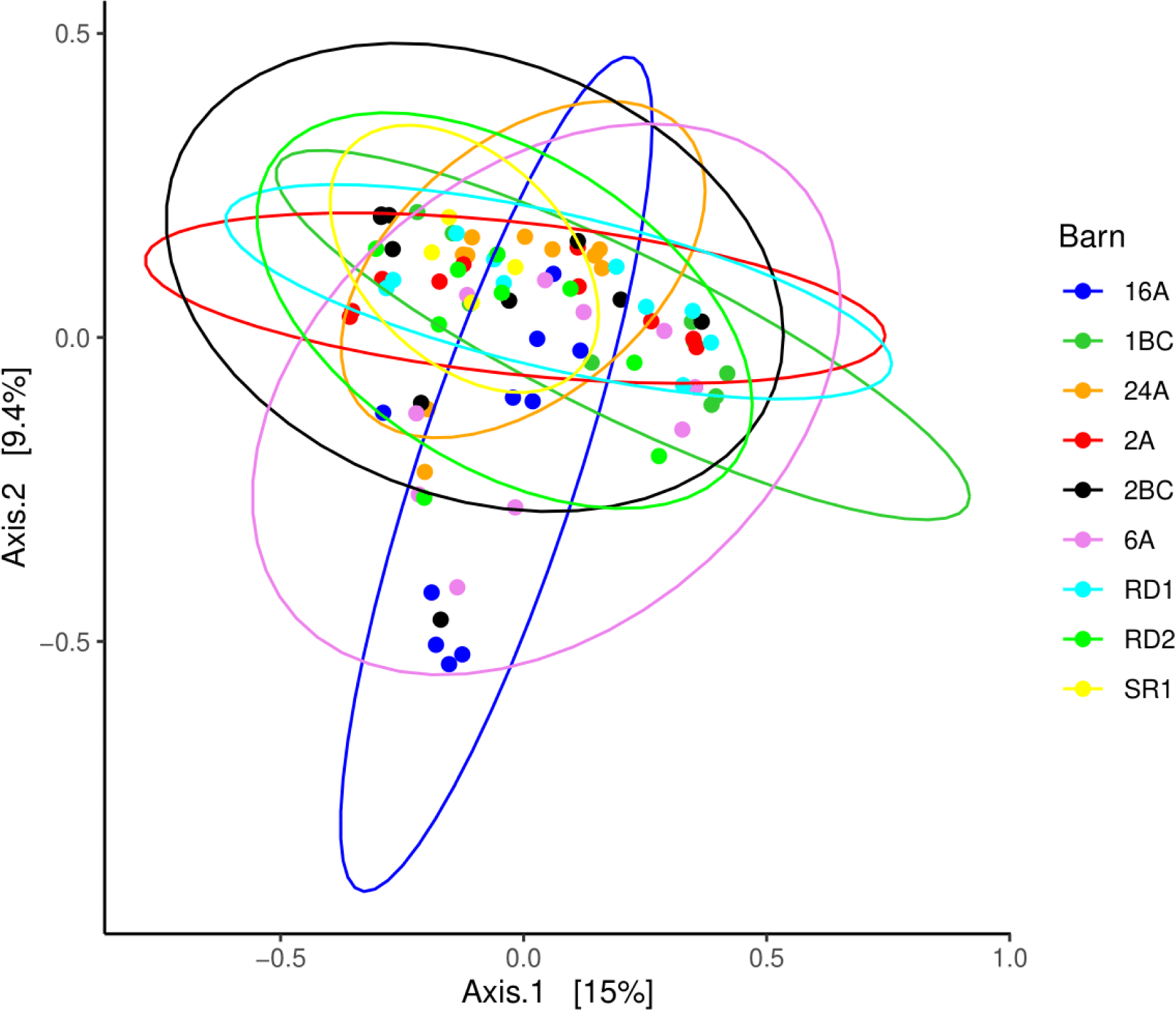
The microbial community structure was different among the domestic pig farms (Adonis P = 0.001, R^2^ = 0.203; beta-dispersion *P* = 0.96) as measured based on Bray– Curtis dissimilarity matrix. The farms in Alberta include 16A, 24A, 2A, 6A, RD1, RD2, and SR1, while those in British Columbia are 1BC and 2BC.

**Supplementary Figure 5:**
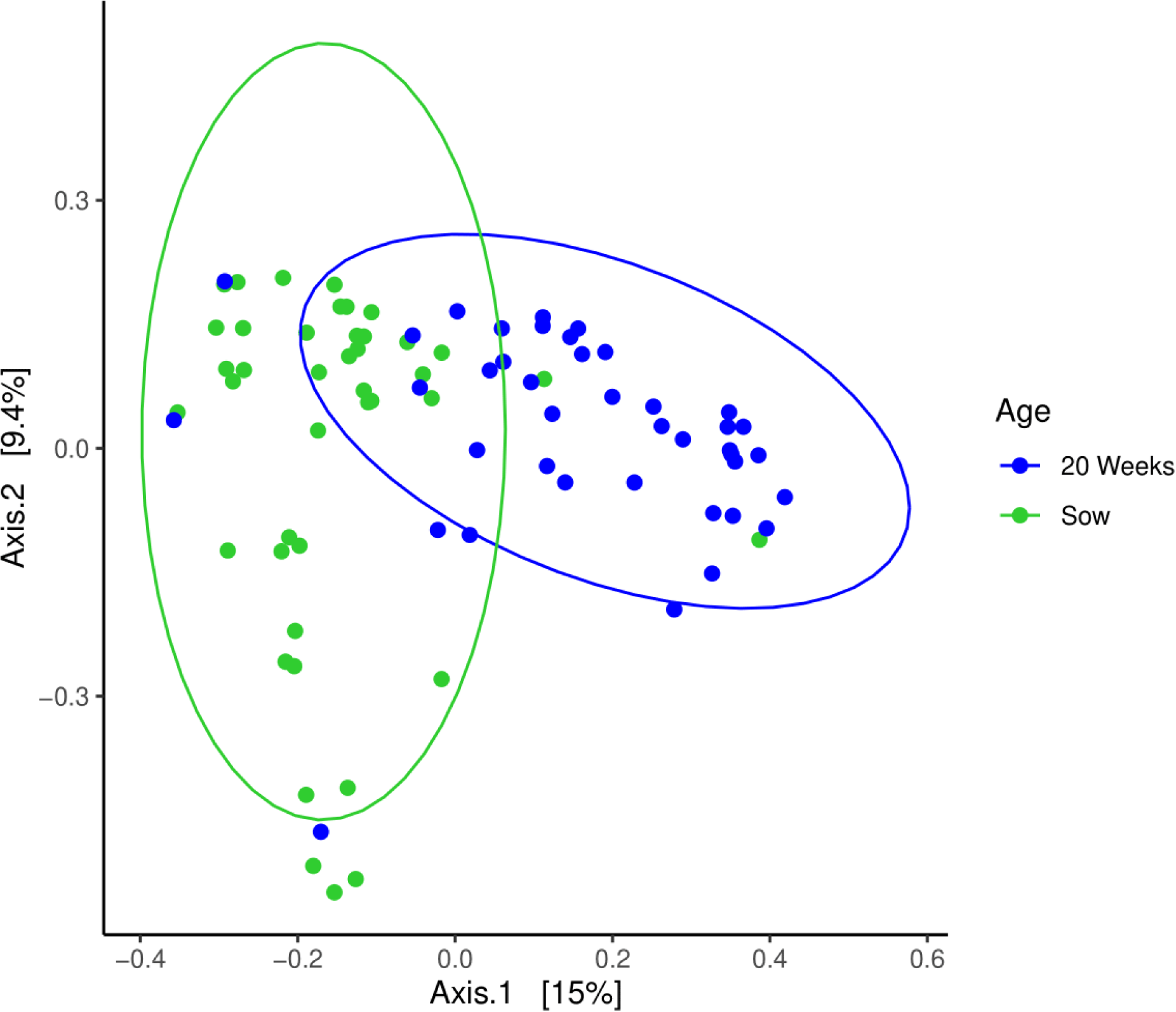
The microbial community structure differed between the Sow and 20-week pigs (Adonis *P* = 0.001, R^2^ = 0.089; beta-dispersion *P* = 0.90) as measured based on the Bray–Curtis dissimilarity matrix.

**Supplementary Figure 6:**
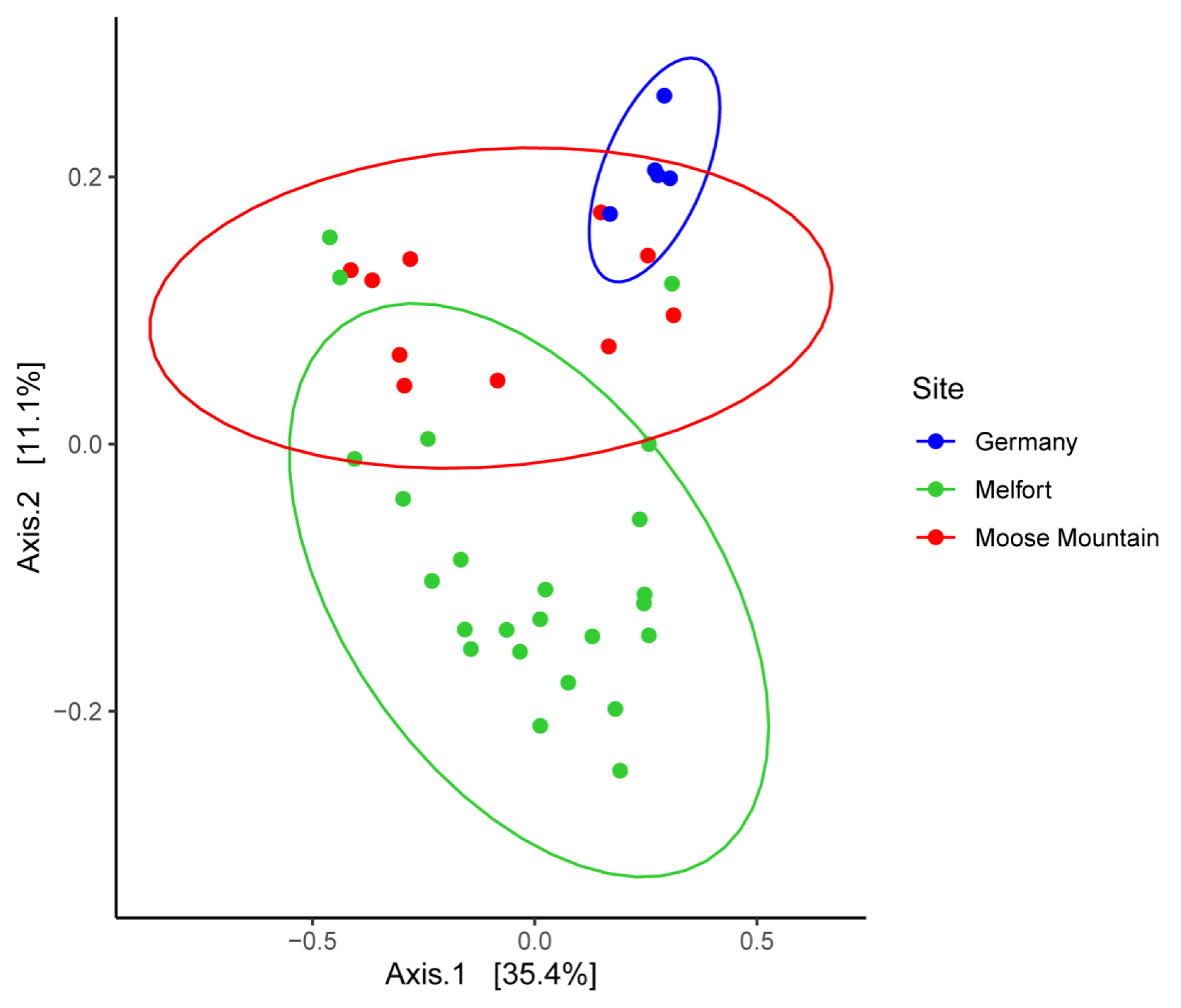
The microbial community structure was different among the wild pig originating locations (Adonis *P* = 0.001, R^2^ = 0.257; beta-dispersion *P* = 0.31) as measured based on Bray–Curtis dissimilarity matrix.

